# A Rational Information Gathering Account of Infant Habituation

**DOI:** 10.1101/2025.01.09.629775

**Authors:** Gili Karni, Marcelo G. Mattar, Lauren Emberson, Nathaniel D. Daw

## Abstract

Gaze is one of the primary experimental measures for studying cognitive development, especially in preverbal infants. However, the field is only beginning to develop a principled explanatory framework for making sense of the various factors affecting gaze. We approach this issue by addressing infant gaze from first principles, using rational information gathering. In particular, we revisit the influential descriptive account of Hunter and Ames (1988) (H&A), which posits a set of regularities argued to govern how gaze preference for a stimulus changes with experience and other factors. When the H&A’s model is reconsidered from the perspective of rational information gathering (as recently also proposed by other authors), one feature of the model emerges as surprising: that preference for a stimulus is not monotonic with exposure. This claim, which has at least some empirical support, is in contrast to most statistical measures of informativeness, which strictly decline with experience. We present a normative, computational theory of visual exploration that rationalizes this and other features of the classic account. Our account suggests that H&A’s signature nonmonotonic pattern is a direct manifestation of a ubiquitous principle of the value of information in sequential tasks, other consequences of which have recently been observed in a range of settings including deliberation, exploration, curiosity, and boredom. This is that the value of information gathering, putatively driving gaze, depends on the interplay of a stimulus’ informativeness (called *Gain*, the focus of other rationally motivated accounts) with a second factor (called *Need*) reflecting the estimated chance that information will be used in the future. This computational decomposition draws new connections between infant gaze and other cases of exploration, and offers novel, quantitative interpretations and predictions about the factors that may impact infant exploratory attention.

## Introduction

Visual attention has long been a primary tool in the study of cognitive processing and development, especially in preverbal infants (Fantz & Ordy, 1959). Gaze (looking time), a commonly used measure of visual attention, has been used to examine language, memory, object recognition, and intuitive physics, among many other topics (Damon, Quinn, & Pascalis, 2021; Frank, Vul, & Johnson, 2009; E. K. Johnson & Jusczyk, 2001; Kidd, Piantadosi, & Aslin, 2012; Kosie et al., 2023; Rose, Feldman, & Jankowski, 2004; Thiessen, Hill, & Saffran, 2005). However, the design of such experiments and the interpretation of the large body of existing results are hampered by the relative lack of a principled explanatory framework or predictive model. Such a framework would ideally serve as a linking hypothesis formally grounding gaze in underlying cognitive processes of interest (Aslin, 2007; Aslin & Fiser, 2005; Poli, Serino, Mars, & Hunnius, 2020). Here we aim to address this gap by revisiting gaze preference — and re-deriving the key features of a canonical descriptive model of it — from first principles, as an instance of rational information gathering. In this, we build upon and extend another recent model (Cao, Raz, Saxe, & Frank, 2022; Raz, Cao, Saxe, & Frank, 2024) that connects gaze and habituation to learning.

Classically, a primary descriptive framework for gaze is the model of Hunter and Ames (1988) (H&A), which aims to systematize a set of factors argued to directionally affect infant visual attention. H&A envisioned that preference between stimuli arises from a hypothetical, covert response to each stimulus, varying with familiarization time. The key feature of the H&A model is that the response has a nonmonotonic, inverted U-shape: as familiarization increases, response initially rises, and then sinks. When comparing some pre-exposed target stimulus to a novel foil, this produces a progression, with exposure, from a transient, early familiarity preference (favoring moderately pre-exposed stimuli over entirely novel ones) to a later novelty preference (disfavoring extensively exposed stimuli). H&A further posit that the overall speed of these dynamics (and hence the trade-off between familiarity and novelty preference, all else equal) is moderated by other factors, notably age and stimulus complexity. Although these general dynamics have not been directly tested in a single study (Kosie et al., 2023), there is much empirical evidence for novelty preferences (Colombo & Bundy, 1983; Fantz, 1964; Rose & Feldman, 1987), and some reports of familiarity preferences (Damon et al. 2021; S. P. Johnson et al. 2009; Lew-Williams and Saffran 2012; Shinskey and Munakata 2005; Sonne, Kingo, and Krøjgaard 2023; Wetherford and Cohen 1973, albeit alongside other studies failing to detect such preferences; Raz et al. 2024).

Although H&A provides an influential descriptive taxonomy for cataloging patterns of directional tendencies, it does not fully serve the ideal of grounding the observable measures in underlying cognitive processes. Its posited U-shaped habituation dynamics are taken as axiomatic, motivated on loose empirical grounds, and not clearly related to functional considerations — particularly cognitive ones like exploration or curiosity. We aim to fill this gap by addressing H&A-style gaze preference dynamics in light of computational models of information gathering, taking the view that gaze is allocated, primarily, to investigate the world. Recent work has used reinforcement learning and statistical decision-theoretic models to investigate a set of common principles motivating information gathering in a range of settings, from covert (deliberation) to overt (exploration in trial-and-error tasks, trivia questions) and has argued that these same underlying principles play out in a range of observable measures including choice behavior, neural recordings, and self-reported boredom and curiosity (Agrawal, Mattar, Cohen, & Daw, 2021; Dubey & Griffiths, 2020; Mattar & Daw, 2018). Here, we build on these theories to argue that the same basic model provides a new, rational underpinning for H&A and the data it describes. In turn, this account proposes a concrete, formal grounding for the abstract, covert variables at the center of the H&A program, which may help to guide, constrain, and interpret experimental work aimed at testing the theory (Kosie et al., 2023).

From a pure information gathering perspective, H&A’s model has one particularly puzzling feature: the signature nonmonotonic form of its habituation function and the resulting transient familiarity preference. In general, statistical informativeness (as measured by changes in uncertainty, entropy, or sampling error, and formalized as the main engine driving gaze in the recent model of Cao et al. 2022) declines monotonically with repeated samples, meaning that on average each new sample resolves less uncertainty than its predecessors. Why, then, should preference for a stimulus, at least according to H&A, increase with exposure, before decreasing?

Here we suggest that this effect can be understood as a manifestation of a feature common to decision-theoretic accounts of many other exploratory domains: While informativeness generally declines strictly with experience, the *value* of information (e.g., for increasing future reward gathering) does not. In particular, in most sequential decision tasks, information about some situation or stimulus is only valuable to the extent that stimulus is likely to be encountered in the future (Anderson & Schooler, 1991; Mattar & Daw, 2018). This common feature of value-of-information calculations has different consequences in different domains: in deliberation, it means we should focus consideration on imminent decisions (Mattar & Daw, 2018); in memory, we should retain the items most likely to be retrieved again (Anderson & Schooler, 1991); in trial-and-error “bandit” tasks, it implies that the value of exploration is modulated by the number of choices left to make (Agrawal et al., 2021; Gittins, 1979; Wilson, Geana, White, Ludvig, & Cohen, 2014); and similar considerations may explain why curiosity and boredom are also impacted by expectations about future events (Agrawal et al., 2021; Dubey & Griffiths, 2020). In the case of gaze, transient familiarity preferences, then, may reflect the expectation that frequently encountered stimuli are likely to be encountered still more in the future.

In what follows, we extend the value of information approach from Mattar and Daw (2018; Agrawal et al. 2021) to build a novel, computational foundation for gaze dynamics and in particular to rationalize the functions proposed by Hunter and Ames (1988). We then use the model to examine current empirical findings and to put forward new predictions for how visual exploratory behavior may vary between situations and individuals, and to clarify the underlying mechanisms driving it.

## Results

### A *need* and *gain* model for visual preference

To clarify the class of tasks we aim to model, Figure 1 illustrates a generic visual preference task. In such tasks, infants are repeatedly exposed to one (or more) stimuli during a familiarization phase. Looking behavior — measured either during familiarization or in later test trials — is used to infer some internal state via the preference for a stimulus vs. other (controlled or implicit) alternatives. These looking patterns are typically measured via time spent looking at a target stimulus, either before looking away or proportionally relative to alternatives. The stimulus display can take different forms. In the version illustrated here (which most straightforwardly instantiates our model), the task is a progression of two-alternative comparisons between a series of novel foils and a repeated stimulus, which gradually becomes familiarized. The central stylized claim of H&A, applied to this task, is that the familiarized stimulus elicits covert arousal that increases and then decreases over repetitions, and is reflected in preferential looking toward it early in training, then toward the novel alternative later. However, our model and its core logic can also be applied to other task variants (as detailed in Methods) such as examining the time to disengage from a single object presented against an environmental baseline.

**Figure 1:**
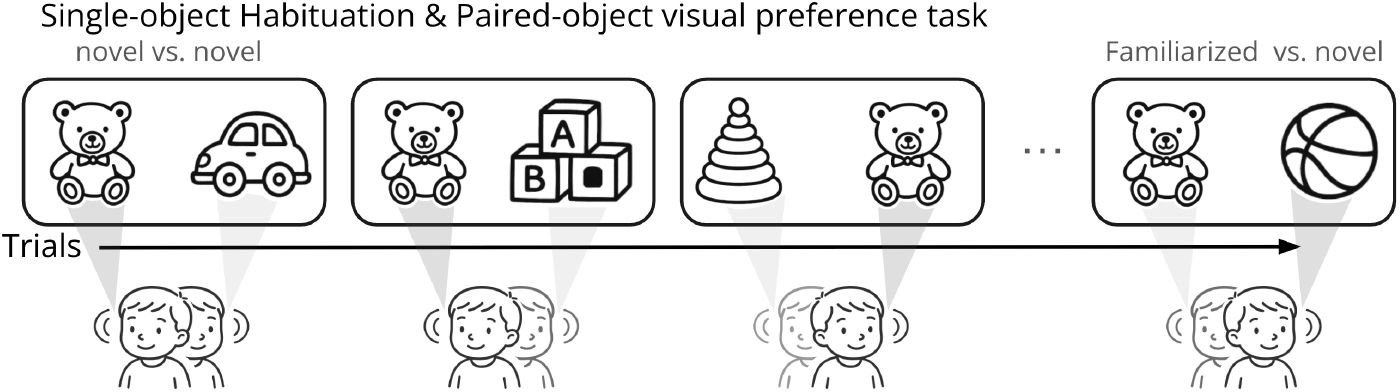
Schematic depiction of a generic visual preference study: Infants are familiarized with a single object through repeated exposure in a paired-object displays showing the familiarized stimulus alongside a novel stimulus. Looking behavior during these paired comparisons is used to infer internal states such as recognition, memory, or preference based on the proportion of time spent looking at each object.

Our aim is to model the dynamics of infant visual habituation by adopting models of optimal information gathering. In particular, this type of model quantifies the value of information-gathering actions in terms of the expected future reward they may enable. Even actions like eye movements that do not directly obtain primary reward may nevertheless ultimately increase rewards gathered, if the information changes decisions the agent later makes in a way that allows them to obtain more reward. For instance, in multi-step trial-and-error reward-gathering tasks, both exploratory choices (Agrawal et al., 2021; Gittins, 1979; White et al., 2019) and steps of planning (Mattar & Daw, 2018) have value in units of reward that can be quantified in terms of how they increase the expected reward earned through later choices. Here we apply this same notion of value of information to infant visual fixations.

To do so, we adopt the mathematical decomposition proposed by Mattar and Daw (2018; Agrawal et al. 2021), who show that in a generic class of sequential decision tasks known as Markov Decision Problems, the value of obtaining information about some “state” (e.g., stimulus) can, suggestively, be decomposed into the product of two terms, which they call *need* and *gain*.

*Gain(s)* captures the increment in cumulative expected future reward, based on the change in the decision that would be made each time state *s* is encountered again: i.e., the return of the better updated choice after obtaining the information, compared to the (worse) choice that would have been made otherwise. But in sequential tasks of this sort, these rewards are only realized if the state where that decision is made is visited in the future; *gain(s)* is thus multiplied by *need(s)*, formally the expected future occupancy of *s* (the expected, time-discounted number of future encounters with *s*). It is then proposed that exploratory actions for each particular option *s* be evaluated by computing (perhaps approximately) its *need* and *gain*, and selected between or prioritized on this basis. (This is also known as “greedily optimal control,” since the proposed prioritization is “greedy” in the sense of not taking account, at each step, of how exploratory actions may also change the value of other subsequent exploratory actions.)

Importantly, experiments on infant visual fixation and habituation like that in Figure 1 do not fully realize the type of decision task analyzed by Mattar and Daw. Although such looking tasks contain decisions about information gathering (i.e., which stimulus to fixate on each trial), they contain no explicit rewards, and so the information subjects obtain is not objectively beneficial in an experimentally controlled way, as it would be in a task like choice between bandits. Thus, in mapping the *need × gain* distinction onto these tasks, only the *need* term is experimentally controlled, via the series of stimulus encounters. For the value of the information, captured by the *gain* term, our model assumes that subjects approximate the value of information as proportional to the amount of information gained (via the expected KL divergence between beliefs, before vs. after sampling). That is, we assume that in choosing which stimuli to fixate, subjects behave as though finding out about some stimulus’ properties will ultimately pay off (incur *gain*) each time the stimulus is encountered again (*need*), even though such rewards are not formally instantiated in the experiments. Heuristics of this sort are standard in exploration (Kaelbling, 1993), Bayesian experimental design (Lindley, 1956), active learning (MacKay, 1992), and cognitive models of curiosity and sampling behavior (Parpart, Schulz, Speekenbrink, & Love, 2017)

More specifically, we assume (similar to Cao et al. 2022; Raz et al. 2024) that infants act as greedy ideal observers encountering a series of stimuli *s* drawn independently (IID) from some initially unknown multinomial distribution with probabilities *p*. The goal of fixating is statistical sampling: each stimulus also has some initially unknown hidden variable(s) associated with it, which can be noisily sampled (or not) by fixating it when encountered. Finally — since we lack a full account of how information gained via sampling will drive later decisions for experimentally controlled rewards — we instead assume that this estimation is itself the goal, i.e., that *gain* for this sampling is taken as ultimately proportional, at each encounter, to the information gained about the hidden quantity by sampling (Kaelbling, 1993).

Accordingly, let us associate the value of fixating some stimulus *S*_*t*_ on some trial *t* with the product of two terms:

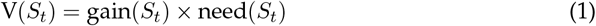

Even before elaborating the detailed specifics of *gain* and *need* (see Methods), we can use slightly simplified proxies for these terms to unpack the intuition why this framework qualitatively reproduces the core dynamics proposed by H&A. The main insight is that these two components change in opposite ways as a stimulus is repeatedly encountered. First, *gain* reflects the benefit of learning more about a stimulus. Because repeated sampling leads to more confident estimates, the marginal gain of each additional observation decreases. That is, the model habituates: the more we know about a stimulus, the less there is to learn (Cao et al., 2022). For example, if a random variable is sampled *n*_*s*_ times, the variance of the sample mean decreases as 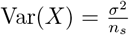. Consequently, the standard deviation of the sample mean shrinks proportionally to 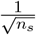, consistent with the law of large numbers. (In the full model, we use a more elaborate Bayesian computation that reflects the same basic logic; see Methods.)

If this were the whole story, then repeatedly sampling a stimulus would have strictly diminishing returns (and preference would always decline with exposure); however, *need* has opposite effects, since expectations about how likely a stimulus is to be encountered in the future depend on how frequently it has been encountered previously. Intuitively, the more often a stimulus has appeared, the more the model expects to see it again. To illustrate: imagine repeatedly rolling a die, where each face represents a different stimulus. The true probabilities of the die’s faces are unknown, but after each roll, we can keep track of how often each face has come up. Starting from a prior assumption that all stimuli are equally likely, each new observation adjusts these estimates. This is a classic Bayesian update, and in this case corresponds to *Dirichlet inference* — a method for estimating multinomial category probabilities from observed counts.

Formally, in our model, stimuli *S*_*t*_ = *s* are drawn from a multinomial distribution with unknown probabilities *p*_*s*_. With a uniform Dirichlet prior (i.e., assuming all stimuli are equally likely at the outset), the updated estimate of *p*_*s*_ after *n*_*s*_ observations is depicted in equation 2 where *j* indexes all stimuli. Because *need* is defined as the expected discounted number of future encounters with stimulus *s*, it grows proportionally with *p*_*s*_, and thus increases as *n*_*s*_ increases. (Again, in the full model we use a version of this computation in which the number of sides of the die is not known ahead of time, but which otherwise has the same properties; see Methods.)

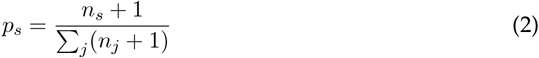

Finally, to connect these quantities to H&A’s thought experiments, we consider a two-stimulus forced choice (e.g. Figure 1). Following H&A, we define preference as the difference between the values of two alternatives, and specifically, novelty preference as the difference between a novel baseline stimulus *s*_0_ encountered *n*_*s*_0 = 0 times and a familiarized stimulus *s* encountered *n*_*s*_ times:

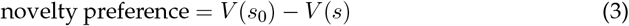

### Novelty and familiarity preferences in the model

Figure 2 shows how the product of *need* and *gain* recapitulates H&A’s classic U-shaped dynamics. With repeated exposures to some stimulus, *need* increases and saturates, while *gain* decreases monotonically to zero. Their product, the value of sampling the familiarized item, rises then falls, mimicking H&A’s proposed U-shaped function. Finally, novelty preference is the difference between the (constant) value of a novel baseline item and the varying value of a progressively familiarized item, resulting in an inverted U-shape function in which the familiar stimulus is first preferred, then the novel one.

**Figure 2:**
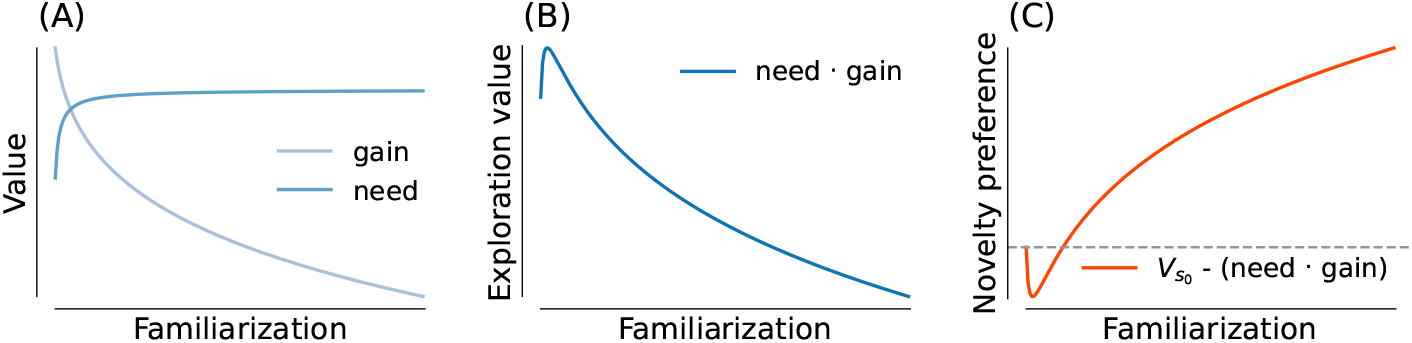
A *need · gain* model of novelty preference. (A) *need* and *gain* components, (B) The value of information through familiarization - guiding exploration, (C) Novelty preference as the difference between a constant value (*V*_*s*_0) of a novel item to the varying one of a familiarized item

While several empirical results support this theoretical model (Balaban & Waxman, 1997; Bogartz, Shinskey, & Schilling, 2000; Kinney & Kagan, 1976; Perone & Spencer, 2014; Roder, Bushnell, & Sasseville, 2000; Rose et al., 2004; Rose, Gottfried, Melloy-Carminar, & Bridger, 1982; Shinskey & Munakata, 2005), we focus on three experiments where results reflect the predicted nonmonotic preference dynamics in different ways. Perhaps the most direct echo of these dynamics is the study of Kidd et al. (2012). In their first experiment, the authors measured infants’ visual attention to target stimuli that appeared, or not, in a sequence with varying levels of appearance frequency versus a blank background stimulus (Figure 3A). Infants were least likely to look away from rare or common stimuli, and more likely to disengage when the appearance frequency was intermediate (Figure 3B). Since stimuli that appear more frequently in the fixed-length series are consequently encountered more times, we can view this result as reflecting a range of familiarization levels analogous to the nonmonotonic preference function from Figure 2. Accordingly, our model captures the result (Figure 3C). (Since this is a one-stimulus experiment, we stylize it, for each probability level, as choice between a target stimulus familiarized a number of times proportional to the appearance frequency, and a fixed baseline value reflecting the rest of the environment.)

**Figure 3:**
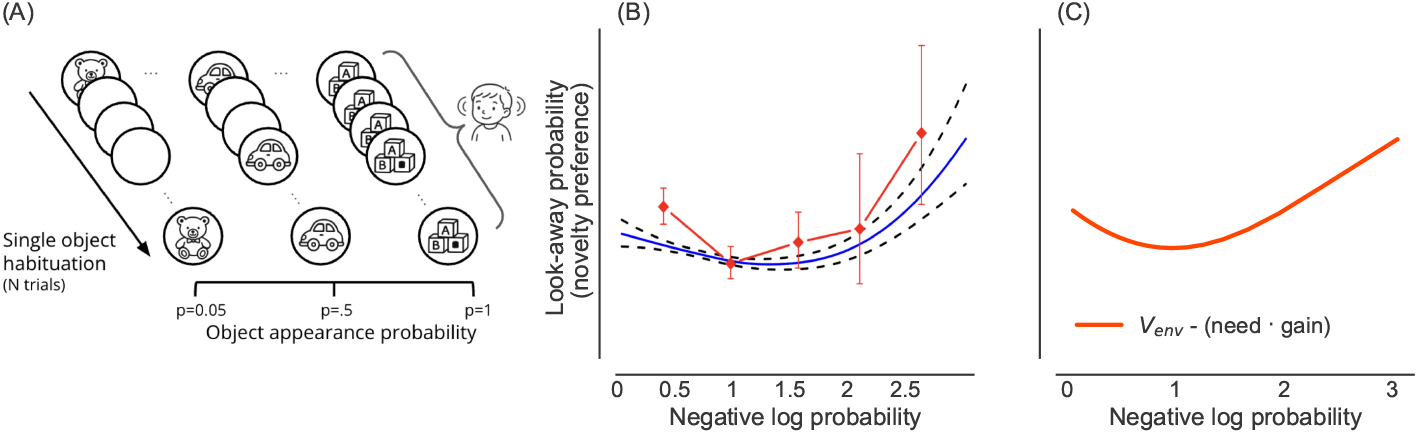
Our model explains empirical results, showing that look-away probability has a U-shaped curve as a function of stimulus appearance frequency. (A) Schematic depiction of the experimental procedures from Kidd et al. (2012): On each trial, infants were shown a single object on a blank background. Each object appeared with a fixed probability across the session. Infants’ looking behavior (i.e., whether they disengaged) was measured on every trial. Analyses compared looking time across appearance probabilities to assess sensitivity to stimulus predictability. (B) Empirical results depicting a U-shape curve of looking away probability as a function of negative log probability. Mean data, with CI, in red (Kidd et al., 2012) (C) Look away probability resulting from the product of *need* and *gain*

An additional demonstration comes from Roder et al. (2000), who tracked novelty preference dynamics across a continuous series of two-alternative choice trials more similar to our stylized experiment in Figure 1. In their task, infants were first shown a brief familiarization trial with a single stimulus, then presented with a series of paired-choice trials where that familiar stimulus was repeatedly paired with different novel stimuli from the same category. In order to facilitate pooling across infants, the data were analyzed relative to the trial when each infant began to show consistent novelty preference, with the trials preceding this further grouped by cumulative stimulus exposure time. The results revealed that infants initially showed familiarity preference during early processing, which gradually transitioned to novelty preference as processing continued (Figure 4B). Since in this trial-based familiarization design, cumulative exposure time accrues over stimulus encounters, it will increase monotonically in the number of encountners and this U-shaped trajectory across processing phases again mirrors our model’s prediction that exploratory value depends on the interaction of decreasing *gain* and increasing *need* with encounters. This is true whether exposure is operationalized via sampling frequency (as in Kidd et al. (2012)) or processing time accumulated over multiple encounters (as in Roder et al. (2000)) (Figure 4C).

**Figure 4:**
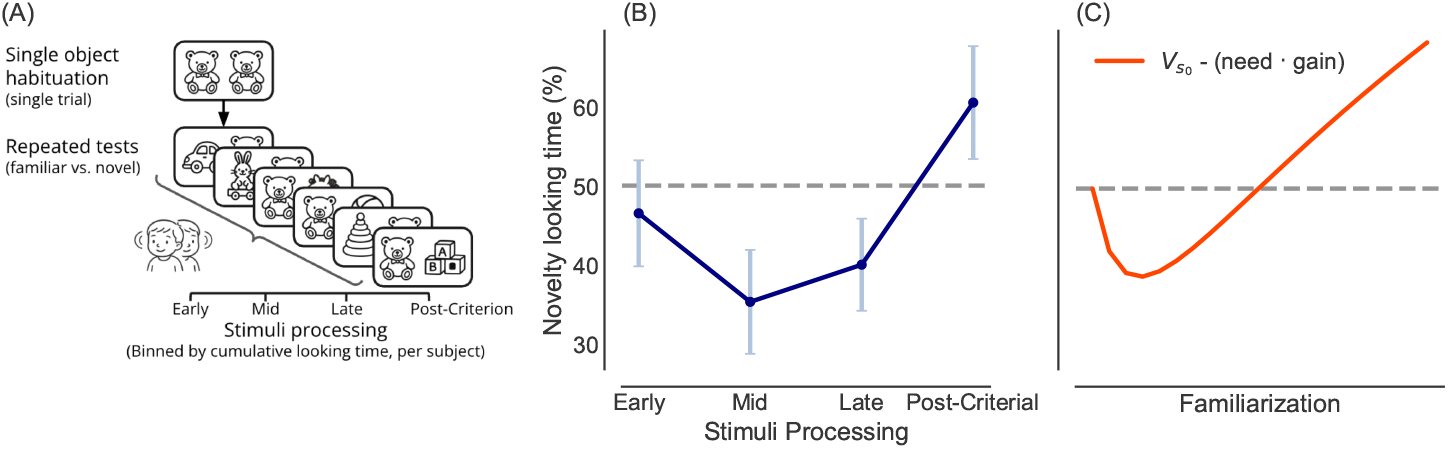
Our model captures the dynamic shift from familiarity to novelty preference during stimulus processing. (A) Experimental schematic from Roder et al. (2000) – Infants were shown a brief familiarization trial, then a continuous series of choice trials pairing the familiar stimulus with different novel stimuli. Preference dynamics were tracked by analyzing looking behavior in trials preceding the onset of consistent novelty preference. (B) Empirical results showing initial familiarity preference transitioning to novelty preference across early, mid, and late processing periods. (C) Model predictions demonstrating how the interaction of decreasing *gain* and increasing *need* produces a similar U-shaped preference trajectory.

A final example from the literature of nonmonotonic habituation — this time in post-exposure delay — suggests that our framework’s principles may extend to memory and forgetting processes. Bahrick, Hernandez-Reif, and Pickens (1997); Bahrick and Pickens (1995); Courage and Howe (1998) habituated infants to a stimulus until they reached a habituation criterion, then tested visual preference for the familiarized stimulus vs. novel ones after varying retention delays ranging up to 3 months. As memory for previously familiar stimuli weakened over longer retention intervals, infants shifted from novelty preference back to familiarity preference (Figure5B). Our model can explain these dynamics if we assume that forgetting, in effect reverses the progression of need and gain that occurs during reversal. That is, as time passes without further exposure, *need* declines (the stimulus becomes less expected) and *gain* increases (due to growing uncertainty), reinflating the exploratory value of previously known stimuli (Figure 5C), with detailed modeling procedures described in the Methods section.

**Figure 5:**
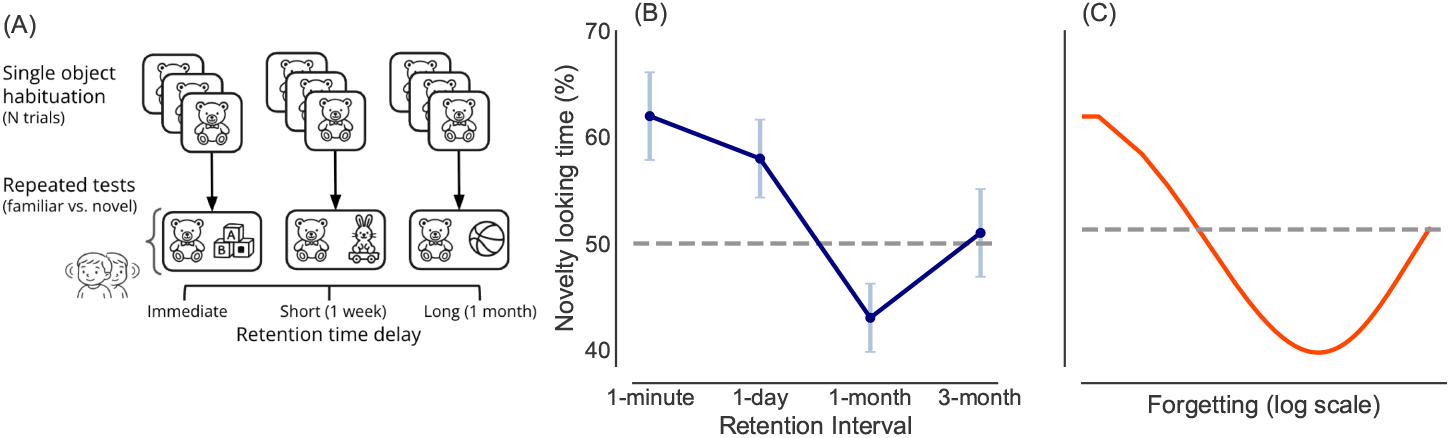
Our framework can explain preference reversals during memory retention, extending beyond learning to forgetting processes. (A) Schematic of experimental procedures from Bahrick et al. (1997); Bahrick and Pickens (1995); Courage and Howe (1998) – Infants were habituated to stimuli, then tested after varying delays (immediate, short – 1 week, long – 1 month) using paired familiar-novel displays. (B) Empirical results showing that as retention intervals increased, infants shifted from novelty preference back to familiarity preference, demonstrating memory-dependent preference dynamics. (C) Model predictions capturing this forgetting-induced preference reversal, where declining *need* and increasing *gain* over time re-inflate exploratory value for previously familiar stimuli.

### Modulators of habituation

Another aspect of the H&A program was the identification of a few factors (notably age and stimulus complexity) hypothesized to affect the progression of the stimulus familiarization dynamics and thus, all else equal, the predominance of novelty versus familiarity preference in any particular situation. Here we explore how different factors underlying *need* and/or *gain* might, in the current model, affect the dynamics of gaze preference. These factors may capture individual differences, cognitive development (or decline), or experimentally manipulable factors. These factors are free parameters of the model, but, importantly (since the model is itself just a generative description of the statistics of stimuli), they reflect features of events that are interpretable and may be experimentally manipulable.

In our model (following Cao et al. 2022; Goodman, Tenenbaum, Feldman, and Griffiths 2008), stimuli are formalized via binary vectors, each element of which is independently drawn from a binomial distribution. Inference then involves estimating the vector of stimulus probabilities from samples. This process drives *gain*, which is modeled as the expected information gain (KL divergence from prior to posterior) upon receiving each sample.

#### Modulators of *Gain*

We first consider stimulus dimensionality, which we use to formalize the notion of stimulus complexity. *Gain* scales with the length of the stimulus vector, such that a higher dimensionality yields a larger initial value of *gain*. These differences, in turn, drive changes in the exploration dynamics (see Figure 6A). Since waning *gain* drives the shift toward novelty preference, larger stimulus dimensionality results in slower habituation: a slower shift from familiarity to novelty preference (see Figure 6B) as in H&A’s model.

**Figure 6:**
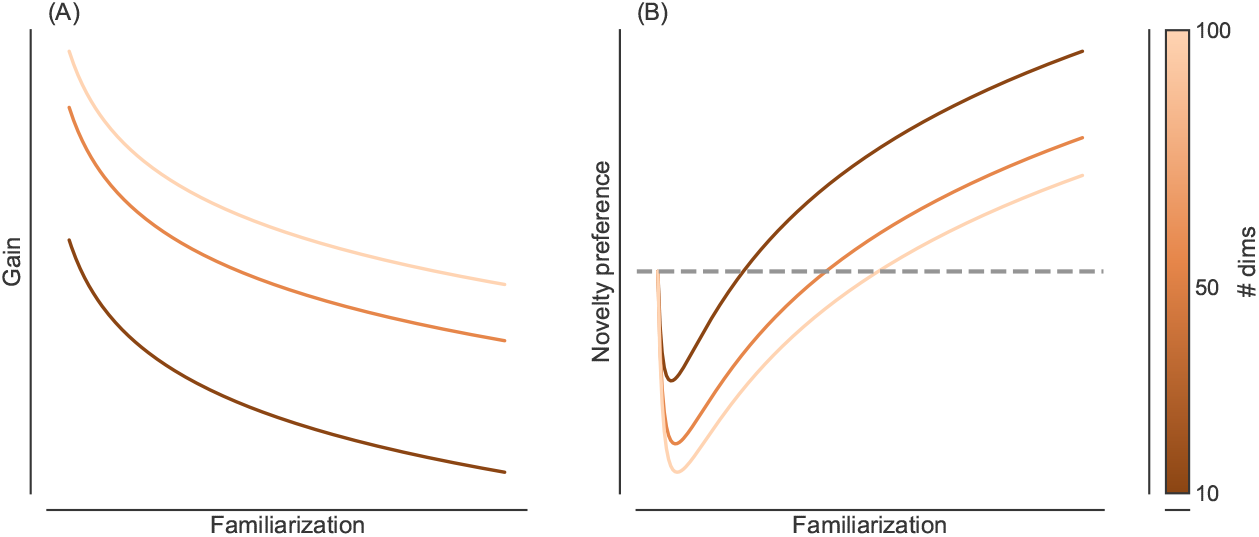
Our model predicts that habituation varies with the number of stimulus dimensions. (A) Larger # of dimensions (f) drives a higher initial *gain* value (B) Larger # of dimensions (f) results in slower habituation

A related parameter is the model’s prior belief distribution over the stimulus probability parameters. Bayesian updating famously involves the precision-weighted trade-off between the prior distribution and the data likelihood. Thus, the speed of Bayesian updating reflects the precision of the prior beliefs, such that more diffuse priors change more nimbly upon sampling data, while more precisely focused priors (whether accurate or inaccurate) update more sluggishly. This means that a stronger prior results in a lower initial gain, and also a flatter, slower decay, compared to a weaker one. Stronger prior precision, in turn, drives flatter, slower habituation, whereas weaker prior precision yields faster, more pronounced habituation (Figure 7B).

**Figure 7:**
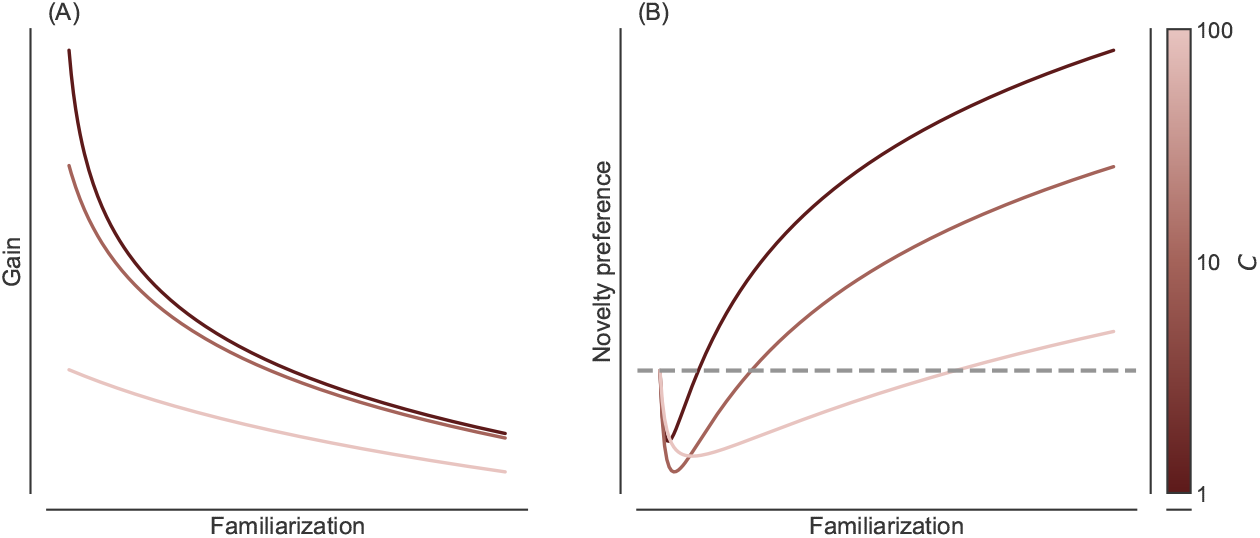
Our model predicts that habituation varies with the precision of prior knowledge. (A) More precise prior belief (higher C) drives a flatter decline of *gain* value, as well as a lower initial value *gain* value (B) More precise prior belief results in flatter and slower habituation

#### Modulators of *Need*

In the model, we assume the infant takes stimuli to be generated according to a distance-dependent Chinese restaurant process. The Chinese restaurant process is a distribution capturing the idea that an unknown number of stimuli will occur with unknown probabilities, like a weighted die with an unknown number of sides. Furthermore, in a distance-dependent Chinese restaurant process, these stimuli are grouped in time; there is some characteristic timescale over which stimuli are likely to repeat.

The key parameter governing the Chinese restaurant process is the concentration or dispersion parameter *α*, which captures how often new stimuli arrive that have never been observed before. A high value of *α* will result in fewer repeated stimuli (in an extreme case, every new stimulus is novel), whereas a low value will yield a denser distribution, where some stimuli repeat many times. Accordingly, a higher dispersion *α* yields *need* that starts lower and saturates more slowly (Figure 8A). Since the saturation of *need* drives the asymptotic dominance of diminishing *gain* (and the shift to novelty preference), these differences drive changes in the exploration dynamics such that higher dispersion *α* results in a slower shift from familiarity to novelty preference (Figure 8B).

**Figure 8:**
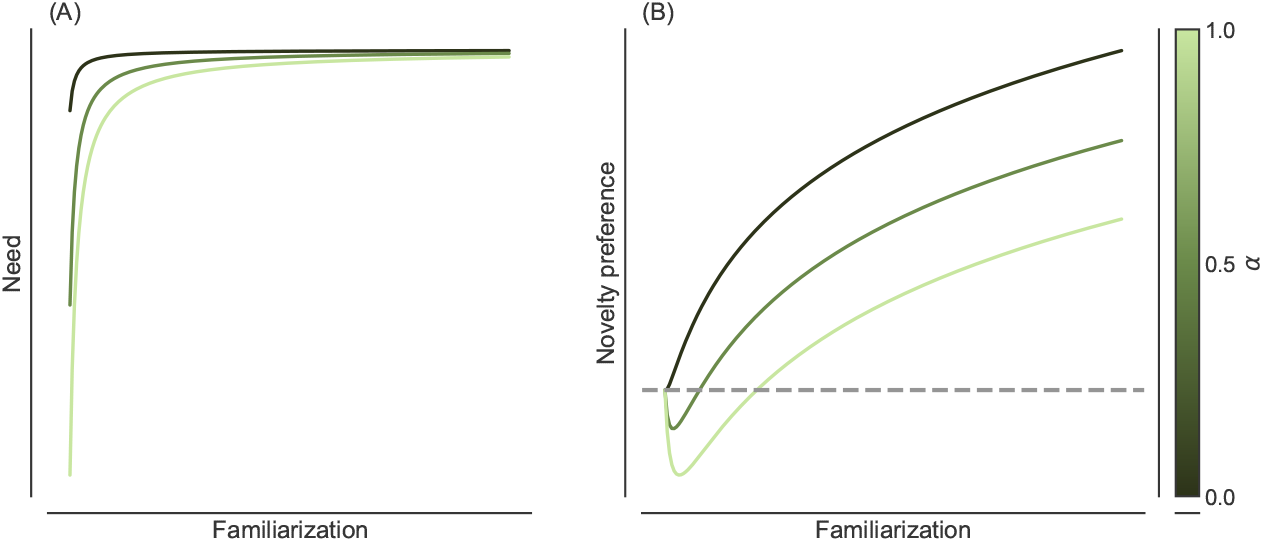
Our model predicts that habituation varies with concentration. (A) Larger concentration (higher *α*) drives a lower initial *need* value (B) Larger concentration results in slower habituation

Finally, a temporal decay parameter *λ* affects the timescale over which stimuli are temporally grouped. A stronger decay permits a smaller window in which stimuli tend to recur, while a weaker decay enlarges this window. Accordingly, stronger decay leads to a faster saturation of *need* with time (Figure 9A). In turn, a stronger decay blunts the impact of *need* yielding more transient familiarity preference and a faster transition to novelty preference (Figure 9B).

**Figure 9:**
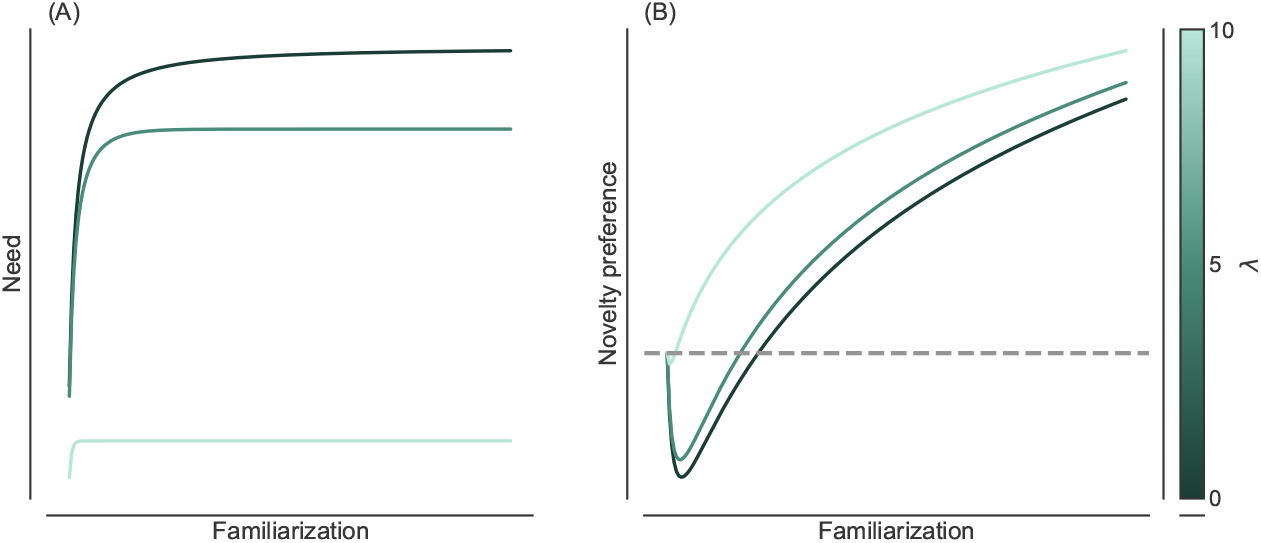
Our model predicts that exploration dynamics vary with temporal decay. (A) Steeper decay (larger *λ*) drives a faster saturation of the *need* value (B) Steeper decay results in faster exploration dynamics.

## Discussion

We propose that significant features of infant exploratory behavior can be captured by a computational model that has previously been used to explain information gathering in other settings in psychology and neuroscience. The model offers a novel rational basis for a predominant account of gaze in developmental psychology, which was not previously expressed in formal terms. A key aspect of the model is that it explains the putatively nonmonotonic preference for stimuli over exposures as the result of the product of two factors: *need* and *gain*, which provide a new theoretical understanding of the rational basis of these phenomena and set the stage for further manipulations testing the theory.

Our model demonstrates the importance of adding *need* to classic information gain considerations to understand the putative nonmonotonicity of exploration and novelty preference. The idea of a decomposing value of information as *need × gain* is not unique to this setting — indeed a major merit of the current proposal is that it provides a unified perspective with other domains including memory (Anderson & Schooler, 1991), deliberation (Mattar & Daw, 2018), exploration (Agrawal et al., 2021), and (in perhaps the closest argument to the current one) curiosity (Dubey & Griffiths, 2020). The use of a common framework suggests that manipulations and measures from these other domains (for instance, manipulating a stimulus’ *need* relative to its competitors while holding absolute experience fixed (Liu, Mattar, Behrens, Daw, & Dolan, 2021) might be brought to bear on infant exploratory behaviors as well.

The *gain* portion of our model draws closely from another recent model of infant gaze, by Cao et al. (2022; Raz et al. 2024), which views moment-to-moment gaze decisions as driven by expected information gain in a similar statistical estimation setting. We adopt as our starting point a similar formalism and rationale, but use a slightly abstracted setting: In particular, we consider sampling as a binary decision at an encounter-by-encounter level, rather than more granular ongoing gaze-or-look away to capture per-stimulus looking times. This simplification is driven by our central goal of augmenting the logic of information gain with the idea that the value of learning about some stimulus or concept is not purely a function of information, but additionally depends on the rate of discrete future encounters with it, i.e. *need* (Anderson & Schooler, 1991).

In our theory, the growth of *need* over multiple encounters drives a transient familiarity preference, as also suggested by H&A. Importantly, this effect is theoretically driven by the target stimulus’ encounter rate relative to other stimuli (Mattar & Daw, 2018) — and not by total exposure time, which is (depending on the situation) only imperfectly correlated with it. We suspect this distinction may contribute to the general sense that H&A’s putative familiarity preference is, empirically, relatively difficult to observe. For instance, the planned ManyBabies 5 experiment (Kosie et al., 2023) modulates familiarization via controlling the length of a single prolonged exposure. In contrast, the U-shaped response measured by Kidd et al. (2012), which we model here, arises as a function of a stimulus disappearing and reappearing repeatedly. Similarly, the U-shaped novelty preference trajectory observed by Roder et al. (2000) may also support this encounter-based interpretation, since although they analyzed data by exposure time, this arose in their design via a series of repeated encounters. We suggest that the distinction between exposure time and encounter rate may help clarify previous and future experimental results. It is less clear, on statistical learning models, to what extent *gain* should depend on the number of encounters versus total looking time, though here we assume the former for consistency and simplicity. This ultimately depends on the assumed noise model by which concept learning is abstracted as noisy sampling of a prototype (Goodman et al., 2008; Raz et al., 2024), i.e., to what extent new information is resampled within versus between encounters.

Our model also suggests two parameters that primarily affect gain, and two that primarily affect *need. Gain* is scaled by stimulus complexity, which has effects consistent with H&A’s proposal that more complex stimuli undergo slower habituation. *Gain* is also affected by the strength of prior information about a stimulus, which might relate to empirical reports of enhanced familiarity preference for common categories like faces (Damon et al., 2021; Houston-Price & Nakai, 2004; Matsuda, Okamoto, Ida, Okanoya, & Myowa-Yamakoshi, 2012).

Importantly, one further aspect of *gain* that simple statistical learning models like ours do not directly capture is that some stimuli are objectively more important, and more valuable to learn about, than others. In common with much other work on novelty preferences and exploration (Brafman & Tennenholtz, 2002; Cao et al., 2022; Kaelbling, 1993), our theory assumes the infant takes learning per se as valuable (at least, learning about stimuli likely to be encountered again), notionally as an approximation to a fuller computation of how particular information helps to obtain later rewards. Nevertheless, an important question for future work is deriving a better formal understanding of how information gain is connected to the objective later reward. Effects of this sort might also contribute to different processing for reward-relevant categories like faces, as well as for other incentives like family members or food.

We also showed how *need* — and in turn, the dynamics of familiarization — is affected by model parameters that capture the (infant’s implied assumptions about the) clustering of stimulus encounters in the environment. These parameters might present interesting candidates for experimental manipulation, insofar as infants might be expected to learn the relevant time horizons by which they expect stimuli to be re-encountered and adapt their *need* computations accordingly. Similar timescale effects are seen in animal and human learning experiments (Piray & Daw, 2021) but have not, to our knowledge, been studied in this setting.

The framework also suggests intriguing connections to memory and forgetting processes. In particular U-shaped novelty-familiarity preference reversal patterns have also been observed in the opposite temporal direction, during forgetting rather than learning. Bahrick et al. (1997); Bahrick and Pickens (1995) and Courage and Howe (1998) showed that as memory for previously familiar stimuli weakens over longer retention intervals, infants shift from novelty preference to familiarity preference. These dynamics can be understood as a temporal mirror of our model: as time passes without further exposure, we expect that need declines (the stimulus becomes less expected) and gain increases (due to growing uncertainty), re-inflating the exploratory value of previously known stimuli. More broadly, the framework generates several testable predictions for future empirical work. If stimulus complexity affects gain as proposed, higher-dimensional stimuli should show prolonged familiarity preference phases. If individual differences in temporal clustering expectations (*α* parameter) exist, they should predict different preference trajectories even with identical exposure sequences. Most directly, manipulating temporal spacing between encounters, while holding total exposure constant, should shift the timing of familiarity-to-novelty transitions, distinguishing encounter-rate effects from exposure-duration effects.

Of course, statistical expectations might also differ between individuals, and could change systematically with an individual’s characteristics, such as age. In this respect, it is worth saying that neither of our *need*-related parameters (those reflecting cluster reliability and the temporal scale) seem likely to coincide with H&A’s proposal that the speed of familiarization dynamics increases with age (Perone & Spencer, 2014; Rose et al., 2004, 1982). For instance, we might plausibly expect older infants to have longer temporal horizons and to form finer stimulus groupings (larger dispersion parameter; Bornstein and Arterberry 2010), but these would actually both promote slower familiarization. Instead, while these effects might indeed exist, we suspect that the overall net effect of age is dominated by gain: i.e., faster, more effective learning, similar to that arising from effectively less complex stimuli. Separate from the explanation for the effect, the possibility that the overall habituation dynamics speed up with age may be another explanation for why familiarity preferences (anyway transient) have been inconsistently detected in experiments. In particular, as exemplified in the recent study of Raz et al. (2024), adults may habituate too quickly for familiarity preferences to be observed, whereas infant gaze is noisy such that experiments may be underpowered for detecting (or, crucially from our perspective, rejecting) a transient effect.

The model presented has a few limitations. First, the underlying model of stimulus or concept learning via binary feature vectors is highly stylized, and surely fails to capture many aspects of how infants learn about the world. We also neglect aspects of stimuli (including value-relevant ones) while considering only familiarization level. Finally, the formal model is abstract and does not speak to the details how quantities like *gain* and *need* might be approximated. Although future work could apply and expand the model to consider these additions, the current model does demonstrate a rational, straightforward account for a previously perplexing phenomenon.

## Methods

### Model definition

The proposed model utilizes Bayesian learning to formalize novelty preference by calculating two quantities: namely *need* and *gain* to determine the exploratory value of observed stimuli. In our discrete temporal setting, the model calculates the exploratory value per target stimulus at every encounter, as a function of the stimuli encountered previously, yielding a metric allowing to contrast multiple stimuli across time. That is, we can now consider the simulated subject’s preference (difference in exploratory value) as akin to the inverse of their “looking away” tendencies as seen in experimental studies (Cao et al., 2022; Kidd et al., 2012; Poli et al., 2020).

### Model of *Need*

*Need* is formalized as distance-dependent Chinese Restaurant Process (dd-CRP) with exponential decay, reflecting a clustering process for non-exchangeable data (i.e., time-dependent), such that stimuli that have been rarely observed are weighted less (Blei and Frazier 2011; Pitman et al. 2002; see equations 4 and 6). This model accounts for the idea that as a stimulus is repeatedly encountered, it becomes increasingly likely that it will be re-encountered in the future; thus (all else equal), it is more worthwhile to learn about. Let *X*_*t*_ denote the assignment of the stimulus presented at time *t*, assuming such assignment occupies the set **K** of categories of familiar stimuli. In a traditional CRP, the probability *p*(*X*_*t*+1_ = *k*|*X*_(1:*t*)_, *α*) is proportional to the number of stimuli previously assigned to category *k*; in our case, this is replaced by a decayed evidence function *ϕ*_*k*_(*t*); with a scaling parameter *α* governing the probability of initiating a new category, that is, a ‘novel’ one. Unlike the traditional CRP, the distance-dependent form addresses the inherent order of appearance invariance, allowing us to consider the sequential nature of our setting.

Here, *α* is the Chinese Restaurant Process dispersion or concentration parameter, reflecting the tendency to assign a stimulus as ‘novel’; the higher it is, the more concentrated the stimuli’s categorizations will be. *λ* is a decay parameter, allowing the model to consider the time difference between stimuli, where a larger value favors more contemporaneous stimuli. Importantly, the *need* model is unconcerned with the form of the stimulus *X*.

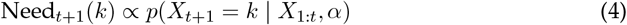

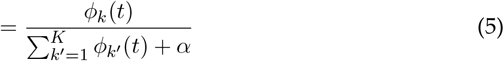

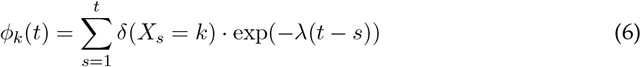

#### Model of *Gain*

The setup for the *gain* model follows a sequential hierarchical Bayesian inference process. We describe the generative model by which stimuli are actually produced in our simulations, and the corresponding inference model (which corresponds to Bayesian inference using the true generative model, except where noted). The agent’s goal is to use a series of per-trial noisy observations to estimate a stimulus vector, ***θ***. Each observation **X**_*t*_ ∈ {0, 1}^*f*^ is a binary feature vector of length *f* (for features) generated independently at each time step *t*. Each feature element *X*_*ti*_ is drawn independently from a Bernoulli distribution with probability given by the corresponding component of the underlying stimulus vector ***θ*** = [*θ*_1_, *θ*_2_, …, *θ*_*f*_]. See Equation 7. This definition ensures that the notation *X*_*t*_ used in the *need* model (Equations 4–6) refers to a time-indexed binary feature vector drawn from this generative process.

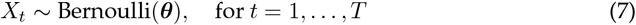

In turn, the vector of probabilities corresponding to a stimulus, ***θ*** is itself generated from a prior distribution representative of some category of stimuli. In particular, a category is given by a vector of means ***µ***, where for generative purposes each element of the stimulus ***θ*** is independently drawn from a beta distribution centered around the corresponding element of the mean ***µ***. In particular, we draw each element of ***θ*** from a distribution *β*(*a, b*), where *a* is equal to a scale parameter *C* = 10 times the corresponding element of *µ* and *b* = *C − a*. (We assume that .1 *≤ µ ≤* .9 in order to ensure that the prior distribution is unimodal.)

For each experiment, we simulate *N* = 200 instances of *S* = 10 stimuli in *K* = 5 categories (where the elements of ***µ*** are each randomly drawn from uniform distributions over [.1,.9]), such that **X**_*k,i,t*_ — the instance at the *t*th trial of the *i*th stimulus in the *k*th category — is drawn from a Bernoulli distribution with probability ***θ***_*k,i*_ corresponding to the *i*th stimulus in the *k*th category (Figure 10 summarizes the generative process).

**Figure 10:**
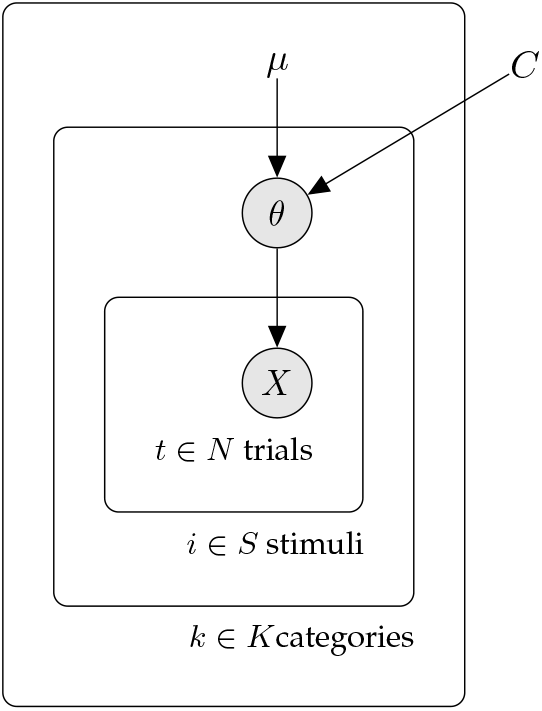
A graphical model of the generative process of the *gain* stimuli. On each trial *t* for stimulus *i* and category *k*, an observation *X* is drawn from a beta distribution with a parameter *θ*, which are determined by category-level means *µ* and non-category-specific constant *C*.

The gain model includes two free parameters. The first is the number of features, *f*, and the second is the strength of the initial prior. For most of the experiments (all those except Figure 7), the prior distribution is taken as maximally uninformative and category-nonspecific: *β*(1, 1) for each element, i.e. uniform. We also (in Figure 7) consider different prior variances, in this case assumed to be category-specific (i.e. centered around the true category means *µ*) but with a range of variances. In this case, we parameterize each element of the prior as *β*(*a, b*) with *a* equaling *C* times the corresponding true category mean from *µ*. Thus we assume the category mean *µ* is known a priori to the subject, and they still aim to infer the stimulus-specific *θ*. However, for the prior variance we consider a range of inferential *C* (e.g, 1, 10, 100, …) both larger and smaller than the generating distribution (*C* = 10), so as to produce prior distributions more or less concentrated around the true category means.

It is important to note that in our model, prior strength and *need* are conceptually and functionally independent. This is because they refer to knowledge at the category vs. exemplar levels. Each stimulus category (e.g., faces or dogs) is associated with a mean *µ* over features (like an average face); each category instance (a particular person’s face) with a noisy sample *θ* around those features; and each encounter with a particular person’s face with a noisy sample **X** around that face’s *θ*. When subjects first encounter a new face, they have a prior that controls their initial uncertainty about *θ*. This in turn reflects their familiarity with, and the dispersion of, the distribution of individual faces *θ* around the category mean *µ*. Need reflects the number of encounters with an individual face, not faces in general. Thus even though a category, like faces, may be familiar (and have a precise prior) individual faces still start with low need when encountered for the first time while different familiarized faces may be encountered more or less frequently (and have higher or lower need).

The model’s belief at trial *t* for stimulus *i* in category *c, p*(***θ***_*k,i,t*_), is given by Equation 8.

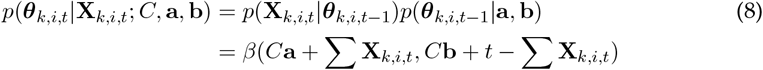

*Gain* is then calculated as the expected information gain (EIG) over the next observation, a metric from Information Theory that has been previously used to measure surprise in cognitive science studies (Cao et al., 2022; Liquin, Callaway, & Lombrozo, 2021; Nelson, McKenzie, Cottrell, & Sejnowski, 2010; Poli et al., 2020; Yin, Chen, Pan, & Tschiatschek, 2021). EIG is approximated as the Kullback-Leibler (KL) divergence between the estimated probability density of some inferred random variable (***θ***) (true for every *k*th category and *i*th stimulus) before versus after sampling another datum (**X**), in expectation over the set of possible observation 0, 1. In Bayesian terms, EIG sequentially models the distance between a prior distribution (*p*(***θ***)) and the expectations of the posterior distribution (*p*(***θ***|**X**)). Equation 9 depicts the EIG over future observation **X** at time *t* + 1 summed over all features *f*. It depicts that a larger update of one’s belief would yield a large value of *gain*. Note that the *gain* is unbounded in its upper limit, as it can be any nonnegative number. To make it compatible with *need* (units of probability), we log-transformed it.

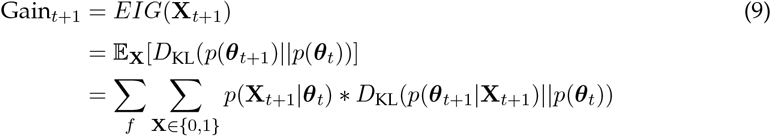

#### Introducing a model of forgetting

We incorporated forgetting mechanisms for both *gain* and *need* components, extending our framework beyond learning to memory retention and enabling the model to capture preference reversals observed in studies with retention delays (Bahrick et al., 1997; Bahrick & Pickens, 1995; Courage & Howe, 1998). **Forgetting Model for *gain***. We modeled information forgetting using a power-law decay function that regresses learned Beta parameters toward a baseline ignorance state (a uniform distribution). Following evidence that memory decay often follows power-law dynamics when aggregated across multiple items (Kahana & Adler, 2002; Murre & Dros, 2015), we implemented forgetting as:

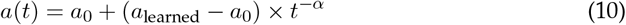

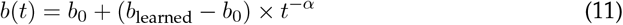

where *a*_learned_ and *b*_learned_ represent the final learned parameters, *a*_0_ = *b*_0_ = 1 represents the uniform prior (ignorance baseline), *t* is time since learning, and *α* = 1.2 controls the decay rate. This formulation captures the psychological intuition that forgetting is most rapid immediately after learning and gradually slows over time (Anderson & Tweney, 1997), contrasting with exponential decay models that predict accelerating information loss. We modeled *need* forgetting through changes in expected stimulus frequency within our distance-dependent Chinese Restaurant Process framework. As *need* represents the predicted frequency of a stimulus category, it increases with repeated observation but decreases as fewer instances are encountered over time. We computed the expected occupancy after observing *N* trials without the target stimulus, where *N* represents the number of trials elapsed since the target stimulus was last encountered. This formulation captures how stimulus categories become less prominent in memory as they are encountered less frequently, with expected occupancy decaying as a function of temporal distance since last exposure (Logan, 1988; Sanborn, Griffiths, & Navarro, 2010). Together, these forgetting mechanisms predict that as time passes without stimulus re-exposure, both uncertainty (*gain*) and categorical expectations (*need*) change in ways that can re-inflate exploratory value for previously familiar stimuli, explaining the preference reversals observed in retention studies.

### Simulations

The model, described above, is then used to draw multiple simulations under different conditions. The *gain* component involves repeated inference of the underlying parameter ***θ*** governing the series of observed binary vectors **X**. Due to the stochastic nature of the model, we present the mean estimate over all simulations.

While manipulating each parameter, we held others constant. Baseline values are set uniformally as prior: *β*(1, 1)), *α* = 1, *γ* = 0.001, *f* = 10, *λ* = 0.001.

### Transparency and Openness

We report all conceptual assumptions, inferential steps, and theoretical commitments in the development of our model. While this is a theory-driven paper, we include illustrative simulations to clarify and demonstrate key predictions. There were no participants, data exclusions, or measured variables. All simulation code and materials are available on https://github.com/karnigili/A-Rational-Information-Gathering-Account-of-Infant-Habituation. This work was not pre-registered, as it does not report confirmatory empirical analyses.

## Acknowledgements

This work was supported by NIMH R01MH135587 (NDD) – part of the CRCNS program, NSERC discovery grant AWD-021239 NSERC 2021 (LLE), and McDonnell Foundation AWD1005451 (LLE).

